# Comparative Transcriptomes of Canine and Human Prostate Cancers Identify Mediators of Castration Resistance

**DOI:** 10.1101/2024.08.18.608482

**Authors:** Marcela Riveros Angel, Bernard Séguin, Christiane V. Löhr, Tomasz M. Beer, John Feliciano, Stephen A. Ramsey, George V. Thomas

## Abstract

Prostate cancer continues to be one of the most lethal cancers in men. While androgen-deprivation therapy is initially effective in treating prostate cancer, most cases of advanced prostate cancer eventually progress to castration-resistant prostate cancer (CRPC), which is incurable. Similarly, the most aggressive form of prostatic carcinoma occurs in dogs that have been castrated. To identify molecular similarities between canine prostate cancer and human CRPC, we performed a comparative analysis of gene expression profiles. Through this transcriptomic analysis, we found that prostatic carcinoma in castrated dogs demonstrate an androgen indifferent phenotype, characterized by low androgen receptor and neuroendocrine associated genes. Notably, we identified two genes, *ISG15* and *AZGP1* that were consistently up- and downregulated, respectively, in both canine prostatic carcinoma and human CRPC. Additionally, we identified several other genes, including *GPX3*, *S100P*, and *IFITM1*, that exhibited similar expression patterns in both species. Protein-protein interaction network analysis demonstrated that these 5 genes were part of a larger network of interferon-induced genes, suggesting that they may act together in signaling pathways that are disrupted in prostate cancer. Accordingly, our findings suggest that the interferon pathway may play a role in the development and progression of CRPC in both dogs and humans and chart a new therapeutic approach.

## 1 Introduction

Most human prostate cancers initially depend on androgen signaling to drive tumor growth.^1^ Thus, for locally advanced or metastatic prostate cancer, androgen deprivation therapy — using antigonadotropins or androgen signaling inhibitors such as abiraterone or enzalutamide —is the backbone of treatment, with an initial response rate of 80–90%.^2,3^ However, most cases that are not cured with local therapies escape medical control and progress to castration-resistant prostate cancer (CRPC) at which point, the disease becomes life-threatening.^4,5^ Despite intensive study, the genomic changes that lead to castration resistance are not fully understood, in part because they are difficult to distinguish from incidental changes due to treatment or disease progression.^5–8^ Comparative oncology studies—which compare molecular features of human cancers with those of related canine cancers under the premise that truly cancer-driving features are more likely to be conserved than incidental features—have revealed molecular mechanisms and led to new directions in cancer detection and treatment for cancers of the bladder, mammary glands, lung, musculoskeletal system, and prostate.^9–17^

Although prostate function in both species is very similar, there are some key morphological and histological differences that are worth noting, as the canine prostate is homogenous in comparison to human prostate which contains four anatomical zones with different glandular arrangement in each zone; the canine prostate is made up 15 lobules, and most of the gland is composed of glandular secretory epithelium. Histologically one of the main characteristics of glands in the canine prostatic tissue is the discontinuity of basal cells, which are seen in a continuous fashion in human prostate and are quintessential for prostatic adenocarcinoma diagnosis as they are usually lost with invasive carcinoma, this is characteristically not seen in canine prostatic cancer.^9,18^ Prostatic carcinoma is rare in dogs, and the routes of metastasis for canine prostatic carcinoma are similar to those of human prostate cancer.^9–12^ Dogs are unique among mammals in their propensity to spontaneously acquire prostatic carcinoma, and they frequently exhibit very aggressive metastatic disease. Research has demonstrated that castration has an impact on the course of prostatic carcinoma, as castrated dogs have a greater likelihood of developing prostatic cancer and experiencing more instances of metastasis compared to uncastrated dogs.^10,19^ A key difference between human and dog prostatic carcinoma is that in dogs, castration (which reduces circulating androgen levels) appears to increase the incidence of spontaneous prostatic carcinoma.^12–14^ We hypothesized that orthologous molecular pathways drive the spontaneous occurrence of prostatic carcinoma in castrated dogs and castration resistance in human prostate cancer, i.e., the canine genes that are transcriptionally up- or down-regulated in castrated dogs with prostatic carcinoma compared to benign canine prostate tissue, are expected to exhibit similarities with the corresponding human orthologs that are up or down-regulated in human CRPC compared to normal human prostate tissue.

To identify these genes, we employed a comparative oncology method. This involved examining the transcriptomes of canine tissue samples and analyzing published RNA seq transcriptome data from humans. By employing a microarray technique on canine prostatic tissue, we identified 33 canine genes that exhibit significantly different levels of gene expression between cancerous and benign prostate tissue. Notably, we found that androgen receptor (AR) regulated genes and neuroendocrine (NE) associated genes were downregulated, suggesting that canine prostatic carcinoma has an androgen indifferent phenotype.

Consequently, we identified five genes (*ISG15*, *AZGP1*, *GPX3*, *S100P*, and *IFITM1*) whose expression profiles indicate consistent dysregulation in both species. We conducted a comprehensive analysis of the protein interaction network of these five genes, examined the rates of somatic mutation co-occurrence, and investigated all canine genes that exhibit differential expression in prostatic carcinoma and demonstrated that these genes are part of a larger network of interferon-induced genes.

This study addresses the knowledge gap that results from most of preclinical models of CRPC relying on immunodeficient mice, cells lines, and genetically modified mice. In contrast, by studying prostate cancer in a species that shares clinical, pathological and tumor microenvironmental features with humans, we have identified a novel interferon signaling network that is dysregulated in CRPC and may reveal treatment targets for both humans and dogs with androgen indifferent CRPC.

## 2 Methods

### 2.1 Tissue samples

Pathology records at the Oregon State University and Colorado State University veterinary hospitals were used to identify fatal cases of prostate cancer in castrated dogs. Thirteen dogs were identified based on pathological diagnosis, clinical information, and postmortem evaluation. Of these, formalin-fixed paraffin-embedded (FFPE) tissue samples were available for tumors from six cases (Figure 1 and Table S1); these were used for the transcriptome study. The FFPE blocks were carefully reviewed before selection, and it was confirmed at the time by histological examination that the selected FFPE blocks contained at least 80% neoplastic cells/tumor fraction. In addition, prostate tissue samples from three dogs that had no diagnosis of prostatic carcinoma and were benign, including both normal prostatic tissue and benign prostatic hyperplasia were used as controls. The three dogs in the control group were non-spayed (not castrated) and used in non-survival cardiac tests at our institution, free of any clinical or pathological indications of prostatic cancer or hyperplasia. Control dogs were young adult males, 1.5-2 years old at the time of tissue collection.

**Figure 1.**
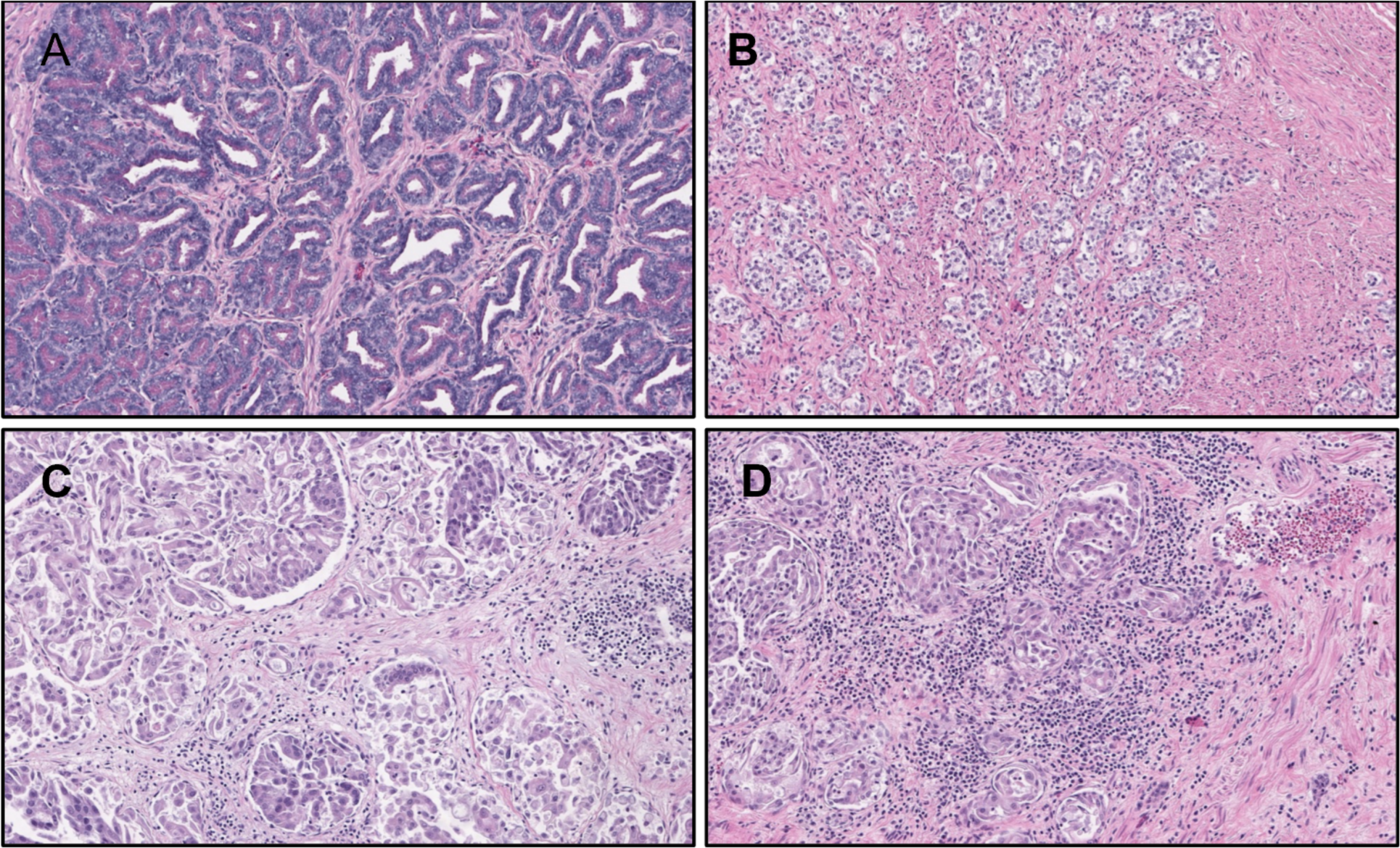
Morphology of benign and neoplastic canine prostate. **A.** Benign prostate tissue (to include normal prostatic tissue and benign prostatic hyperplasia). **C, B and D:** The images illustrate examples of prostatic carcinoma, lobulated and nested clusters of tissue, some forming glands, with invasive characteristics and areas of poorly formed glands and fusion of ductal epithelium (H&E stains, 200x magnification).

### 2.2 Microarray profiling

FFPE canine tissue samples (*n* = 5 from CSU; *n* = 1 from OSU) were sectioned on a Leica microtome at 10 μm thickness. RNA was isolated from tissue sections following the miRNeasy FFPE kit protocol (Qiagen, Redwood City, CA) which includes an on-column DNAse step. RNA yields, as measured on the NanoDrop ONE spectrophotometer (Thermo Fisher, Waltham, MA), were in the range of 1.6–2.8 μg for the small tissue samples and 13.7–66.3 μg for the large samples. Absorbance ratios (260 nm / 280 nm) for the samples were in the range of 1.89 ± 0.04 (s.d.). Samples were assayed on the Agilent Bioanalyzer 2100 using the RNA 6000 Nano Chip on the eukaryotic Total RNA program; DV200 levels for the samples were in the range of 59% ± 17% (s.d.), indicating adequate integrity for microarray profiling. From 9 total RNA samples, labeled target single-stranded complementary DNA was synthesized using the NuGEN FFPE Whole Transcriptome Amplification System (Redwood City, CA) with the Encore labeling protocol. Amplified and labeled cDNA target samples were each hybridized to an Affymetrix GeneChip Canine Gene 1.0ST array (Thermo Fisher, Waltham, MA). Sample-level GeneChip image files (“.dat” files) were processed to probe intensity-level data (“.cel” files) using the Affymetrix Command Console v.3.1.1 software (Thermo Fisher, Waltham, MA).

### 2.3 Bioinformatics

#### 2.3.1 Microarray data analysis

We analyzed probe intensity-level data using the R statistical computing software (ver. 4.0.3) and using R packages from the Bioconductor project (ver. 11) and using R packages installed from the Comprehensive R Archive Network website, as detailed below. To process Affymetrix probe-level intensity data into probeset expression level estimates, we used the packages “oligo” (ver. 1.54.1) and “affycoretools” (ver. 1.62.0) from Bioconductor, with probe annotations from the package pd.cangene.1.0.st (ver. 3.12.0). We used the robust multichip average method of the oligo package for background adjustment, quantile normalization, and probeset summarization.^20^ We annotated probesets using the NetAffx build 36 (build date March 15, 2016) annotation data file for the microarray chip CanGene 1.0 ST (ver. 1; downloaded from the Affymetrix NetAffx Analysis Center website). To eliminate background-level genes, we filtered for probesets that reached an intensity of 16 in at least one sample. We carried out the Principal Coordinates Analysis using the R functions “dist” (with the default Euclidean distance measure) and “cmdscale.” We tested probesets for differential sample-group-average intensities using the Bioconductor package “limma” (ver. 3.44) with empirical Bayes-based variance estimation.^21^ For the comparison to the canine bladder cancer samples, we used microarray data from NCBI GEO (accession number GSE183793), which used the same Affymetrix canine gene ST array therefore enabling direct comparison with our dataset. This dataset consisted of three K9TCC bladder cancer cell line samples and we co-normalized these with our canine prostate and canine prostate tumor samples via probeset quantile normalization (Figure S1).

#### 2.3.2 RNA sequencing data analysis

We obtained previously published RNA-seq transcriptome data from human prostate cancer and normal prostate tissues from the Cancer Genome Atlas (TCGA) prostate adenocarcinoma (PRAD) study using the web-based tool Xena.^22,23^

#### 2.3.3 Protein network analysis

We used the GeneMania web tool, which aggregates information from 889 different published human molecular datasets or databases (with the automatically selected weighting method and with a maximum of 20 resultant added genes) with the following five gene symbols provided as seed genes: *ISG15*, *AZGP1*, *GPX3*, *S100P*, and *IFITM1*, to search for a molecular network community integrating the seed genes.^24^ The resulting network contained 25 genes (each mapped to a specific protein) and five different types of edges: (1) coexpression across various cell or tissue types assayed for gene expression in 377 studies whose datasets have been deposited in the National Center for Biotechnology Information’s Gene Expression Omnibus (GEO) database;^25^ (2) physical interactions reported in any of 432 datasets of protein-protein interactions; (3) predicted physical interactions reported in any of 51 datasets of computationally predicted protein-protein interactions (e.g., based on orthology to interacting protein pairs in other species); (4) gene-gene interactions reported in any of 23 genetic interaction datasets; and (5) co-participation in a biomolecular pathway reported in any of six pathway databases.

#### 2.3.4 Somatic mutation analysis

We used the web-based tool cBioPortal to obtain gene-level data on somatic mutation types and frequencies for human prostate cancer, including co-occurrence frequency data and copy number variation frequency data by variant type (amplification or deletion).^26^

#### 2.3.5 Pathway enrichment analysis

For pathway enrichment analysis, we used the ShinyGO ^27^ tool version 0.77 (accessed on Nov. 2, 2023), which is based on gene annotations from Ensembl Release 104.

## 3 Results

### 3.1 Canine prostatic carcinoma is androgen indifferent

We profiled transcriptomes using RNA recovered from banked FFPE tissue. Specifically, because of RNA integrity limitations due to archival FFPE material, we used a hybridization microarray of oligonucleotide probes targeting 3’ ends of canine genes (instead of RNA sequencing) to measure abundances of 32,391 canine genes in eight samples (five prostatic carcinoma tumor samples and three benign prostate tissue samples), of which 23,389 were above background (see Methods). To investigate the molecular differences between bladder and prostate tissues, as well as between normal prostate and prostatic cancer states, we performed Principal Coordinate Analysis (PCoA) on the collected samples (Figure S1). The first principal coordinate (PCo1) clearly separated bladder tumor samples from both normal and tumor prostate samples, accounting for the largest proportion of variance in the dataset. This pronounced separation along PCo1 suggests fundamental molecular differences between bladder and prostatic cancer cells and provides additional level of support for their separate biology and derivation. Interestingly, while prostate samples showed some overlap, there was a noticeable separation between normal and cancer prostate samples primarily along the second principal coordinate (PCo2). This separation indicates detectable molecular alterations associated with prostate tumorigenesis.

Next, In an unsupervised principal coordinates analysis, the cancer and benign sample groups were separable at the level of the second principal coordinate (Figure S2). For the above-background genes, we analyzed the probe-set intensities for differential expression between sample groups, and identified 33 canine genes that have significantly altered (i.e., FDR < 0.3 and an average expression ratio of at least 1.5 between the two sample groups) transcript abundances in cancer vs. benign prostate (Table 1; Table 2); these comprised seven upregulated genes with higher transcript abundance in cancerous than in benign prostate tissue and 26 downregulated genes with lower abundance in cancerous than in benign tissue.

**Table 1.** Summary of the 33 genes that are differentially expressed (false discovery rate, i.e., FDR < 0.3 and at least a 1.5-fold change) between canine prostatic carcinoma (n = 5) and benign prostate (n = 3) tissue in neutered dogs. Symbol, official gene symbol; Probe-set, the Affymetrix probeset identifier for the target transcriptomic locus; log_2_(C/N), log_2_ cancer/benign mRNA.

| Symbol | Description | Probe-set | $\log_2(C/N)$ | q |
| --- | --- | --- | --- | --- |
| *ACP3* | acid phosphatase, prostate | 14345131 | -3.66 | 7.6E-04 |
| *ANK3* | ankyrin 3, node of Ranvier (ankyrin G) | 14406048 | -2.09 | 4.5E-02 |
| *APOC1* | apolipoprotein C-I | 14264943 | 0.87 | 2.9E-01 |
| *ARHGAP28* | Rho GTPase activating protein 28 | 14432754 | -2.21 | 2.0E-02 |
| *ATP6V0A4* | ATPase, H<sup>+</sup> transporting, lysosomal V0 subunit a4 | 14299605 | -2.21 | 3.7E-02 |
| *AZGP1* | α-2-glycoprotein 1, zinc-binding | 14424840 | -2.24 | 5.8E-02 |
| *BCAS1* | breast carcinoma amplified sequence 1 | 14352265 | 1.41 | 4.6E-02 |
| *CACNA2D1* | Ca channel, voltage-dependent, $\alpha$ 2 / $\delta$ subunit 1 | 14313786 | -4.44 | 1.1E-03 |
| *CLDN10* | claudin 10 | 14342285 | -4.35 | 7.0E-04 |
| *CMTM8* | CKLF-like MARVEL transmembrane domain-containing 8 | 14346402 | -2.83 | 1.1E-02 |
| *CXXC4* | CXXC finger protein 4 | 14388493 | -2.51 | 7.6E-03 |
| *DNASE1L3* | deoxyribonuclease I-like 3 | 14328198 | -6.10 | 5.9E-04 |
| *ELAPOR1* | KIAA1324 ortholog | 14428037 | -2.56 | 7.6E-03 |
| *GPX3* | glutathione peroxidase 3 (plasma) | 14407751 | -3.03 | 7.1E-02 |
| *IFITM1* | interferon-induced transmembrane protein 1-like | 14310666 | 4.11 | 2.4E-01 |
| *IGLL5* | immunoglobulin lambda-like polypeptide 5-like | 14359030 | 2.05 | 3.0E-01 |
| *ISG15* | ubiquitin-like protein ISG15-like | 14411891 | 3.66 | 2.7E-01 |
| *KLK2* | kallikrein-related peptidase 2 | 14264046 | -6.98 | 1.0E-04 |
| *KYNU* | kynureninase | 14317645 | -3.78 | 6.8E-02 |
| *MME* | membrane metallo-endopeptidase | 14345787 | -3.39 | 4.6E-02 |
| *NKX3-1* | NK3 homeobox 1 | 14353801 | -4.30 | 8.6E-04 |
| *PHYH* | phytanoyl-CoA hydroxylase-like | 14319723 | -3.93 | 2.3E-03 |
| *PIP* | prolactin-induced protein | 14301387 | -3.05 | 4.6E-02 |
| *PON3* | paraoxonase 3 | 14292642 | -4.48 | 4.7E-03 |
| *RNF19A* | ring finger protein 19A, E3 ubiquitin protein ligase | 14467680 | 2.11 | 1.9E-01 |
| *S100P* | S100 calcium binding protein P | 14375352 | 3.27 | 2.7E-01 |
| *SLC12A2* | solute carrier family 12 (Na/K-Cl transporters), member 2 | 14274245 | -4.33 | 2.9E-02 |
| *SLC38A11* | solute carrier family 38, member 11 | 14396551 | -2.82 | 1.1E-03 |
| *SPP2* | secreted phosphoprotein 2, 24 kDa | 14354497 | -1.90 | 6.4E-02 |
| *STEAP1* | six transmembrane epithelial antigen of the prostate 1 | 14290426 | -2.39 | 4.6E-02 |
| *STEAP4* | STEAP family member 4 | 14292450 | -1.85 | 9.8E-03 |
| *TMEM213* | transmembrane protein 213 | 14301689 | -1.81 | 4.0E-02 |
| *LYSET* | lysosomal enzyme trafficking factor | 14439109 | -1.71 | 4.5E-02 |

**TABLE 2.** Seven sample group comparisons for transcript abundance analysis in this study.

| Data source | Species | Sample group 1 | Sample group 2 |
| --- | --- | --- | --- |
| this study | dog | PAC | benign prostate |
| TCGA <sup>22</sup> | human | PC | normal prostate |
| Yun et al. Oncotarget <sup>6</sup> | human | advanced PC | PC |
| Yun et al. Oncotarget <sup>6</sup> | human | CRPC | PC |
| Yun et al. Oncotarget <sup>6</sup> | human | CRPC | BPH |
| Yun et al. Oncotarget <sup>6</sup> | human | PC | BPH |
| Stand up to Cancer (SU2C) <sup>34</sup> | human | CRPC (+abiraterone) | CRPC (-abiraterone) |

Plus (+) and (-) symbols indicate CRPC tumors that from abiraterone-treated patients and from patients that had not previously been treated with abiraterone, respectively. Sample group ratios reflect the average of group 1 over the average of group 2.

Three of the genes that are downregulated in canine prostate cancer are known to be androgen-regulated: *KLK2* (encoding prostate-specific arginine esterase)*, NKX3.1* (encoding NK3 homeobox 1), and *ACP3* (encoding prostatic acid phosphatase).^28,29^ *KLK2*’s human ortholog is a paralog of *KLK3,* the human gene that encodes prostate specific antigen (PSA).^10,30–33^ *ACP3’*s human ortholog, PSAP is a human prostate cancer biomarker.^31^ Consistent with the finding that these three androgen-regulated genes have lower transcript abundance in prostate cancer than in benign prostate, we noted a decrease in the expression of the AR gene in the castrated canine prostatic carcinoma, as compared to the benign prostate (Figure 2A); a finding that aligns with the approximately 1.5-fold shift in the distribution of *AR* transcript abundance between human normal prostate and localized prostate cancer based on data from the Cancer Genome Atlas (TCGA) prostate cancer study (Figure 2b).^22^ Conversely, in humans with metastatic CRPC, AR signaling is increased, usually due to amplification or mutation of *AR.*^34^ Overall, these results suggest that increased AR signaling is not common in prostate cancers arising in castrated dogs in contrast to human metastatic CRCP.

**Figure 2.**
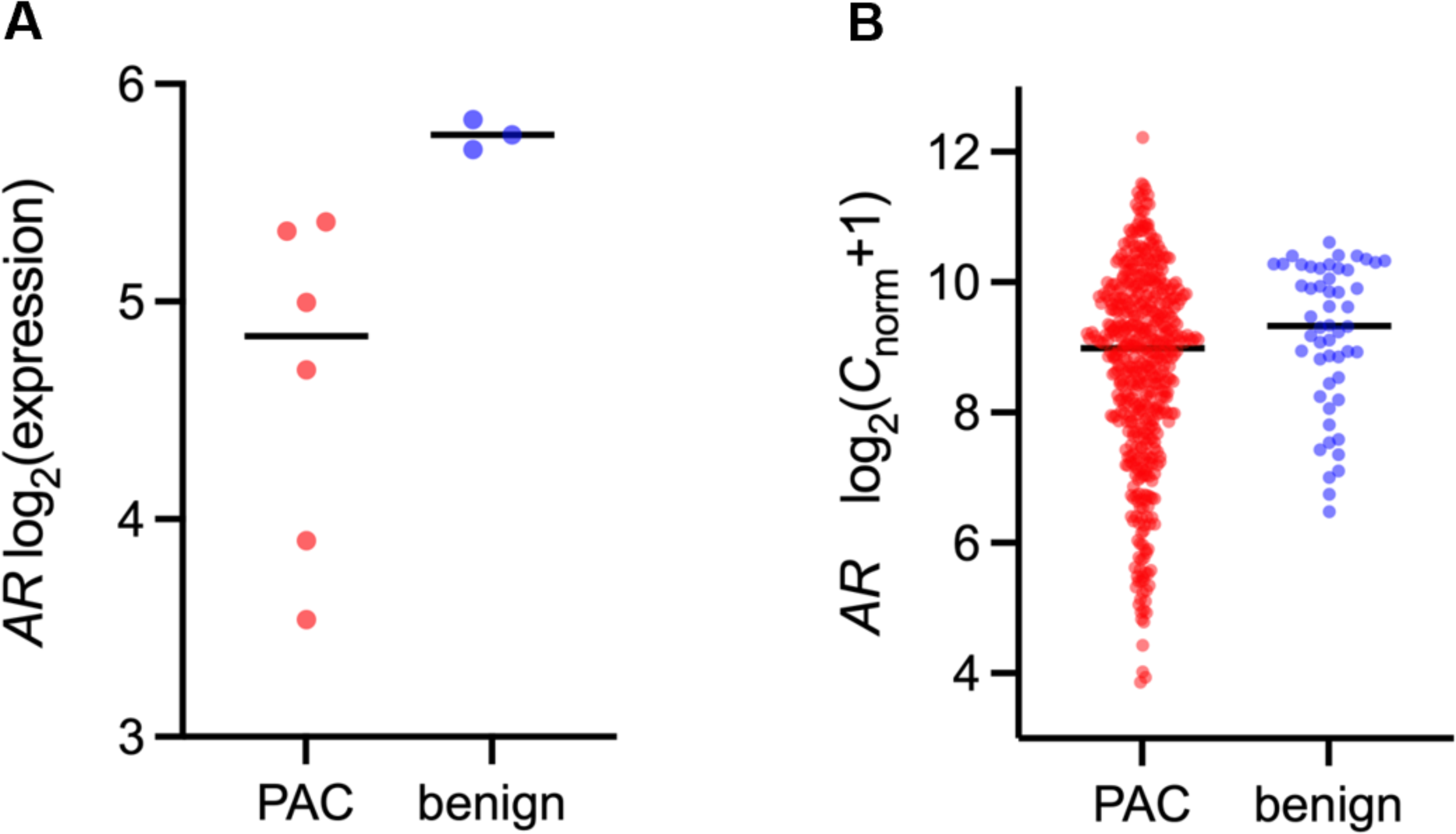
Comparison of Androgen Receptor (AR) transcript abundance in both benign canine tissue vs prostatic carcinoma and benign human tissue vs localized prostate carcinoma. **A.** In canine tissue sample-group-average AR transcript abundance is lower in prostatic carcinoma than in benign prostate tissue. **B.** In human tissue, from TCGA expression data, there is a 1.5-fold difference in the levels of AR transcript abundance between benign human prostate and localized prostate cancer. Marks indicate individual tissue samples and bars indicate median.

Next, we examined the expression of genes associated with the neuroendocrine phenotype: *INSM1*, *RB1*, *TP53*, *CHBA*, *CHGB*, *SYP*, and *SYPL1*. In both benign canine prostate and canine prostatic carcinoma, the absolute, sample-group-averaged probeset intensities for these genes were low (i.e., log_2_ intensity of less than six) (Figure 3). Though we did not have immunohistochemistry markers to characterize neuroendocrine differentiation, the pathology did not demonstrate small cell or neuroendocrine features (Figure 1). Taken together, our findings suggest that prostatic carcinoma in castrated dogs is more aligned with the androgen receptor null, neuroendocrine associated null, i.e. “double-negative” phenotype.

**Figure 3.**
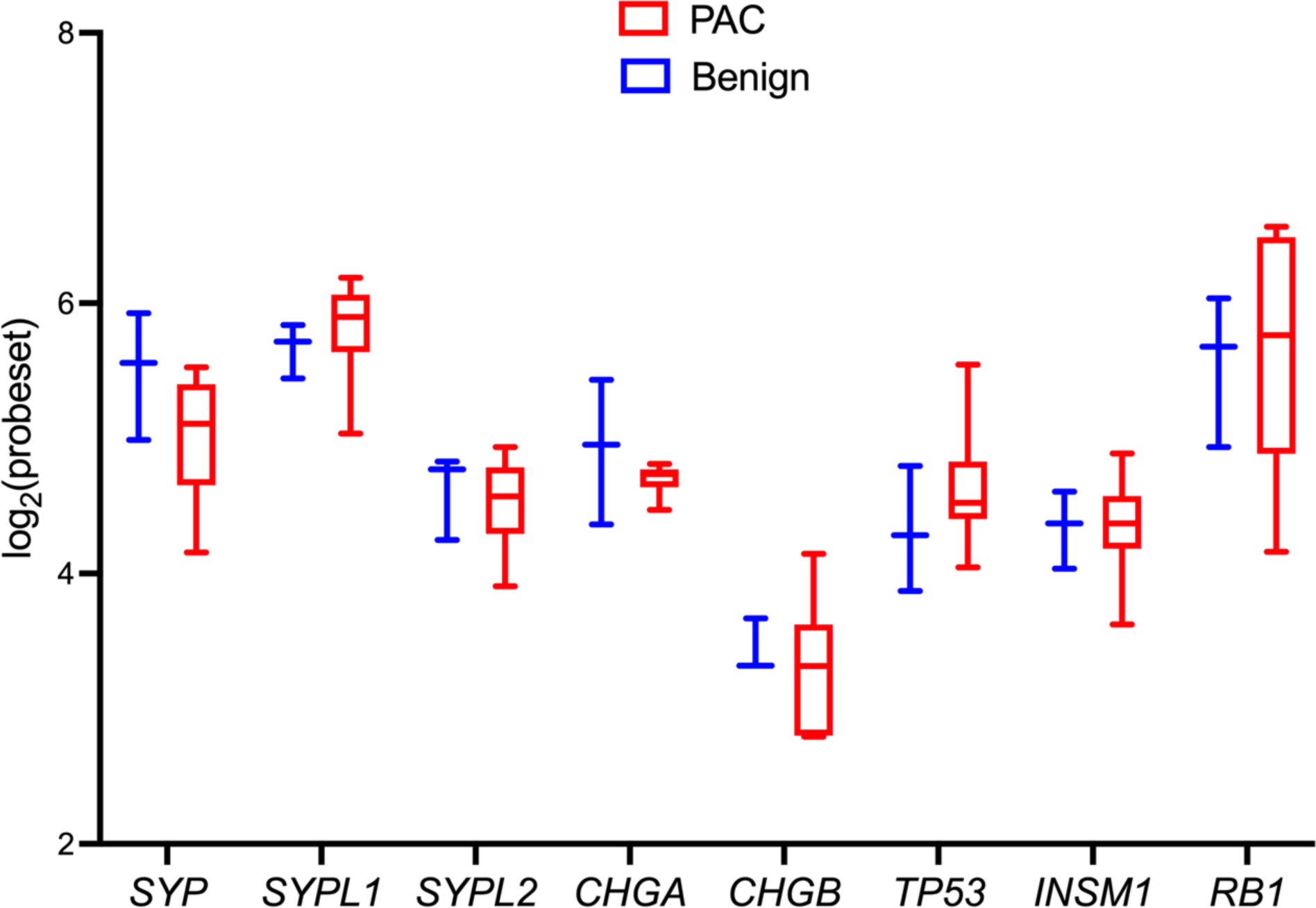
Expression of common genes associated with neuroendocrine carcinomas phenotypes in canine prostatic carcinoma and benign tissue. The gene expression levels of neuroendocrine genes *INSM, RB1, TP53, CHGB, CHGA*, and *SYPL* show no substantial disparity between prostatic carcinoma and benign prostate tissue. These findings, together with the tumor’s histologic characteristics, demonstrate the tumor phenotype is distinct from neuroendocrine prostate carcinoma.

### 3.2 Gene expression reveals commonalities of castration resistant prostate cancer in humans and dogs

Next, to explore the functions of the 33 genes that are differentially expressed in prostatic carcinoma in castrated dogs, we mapped the 33 genes to human orthologs and obtained measurements of the orthologs mRNA abundance ratios between CRPC and localized prostate cancer.^6^ We co-analyzed canine-human ortholog pairs by first examining canine cancer/benign expression ratios and, secondly human CRPC/PAC expression ratios (Figure 4A).

**Figure 4.**
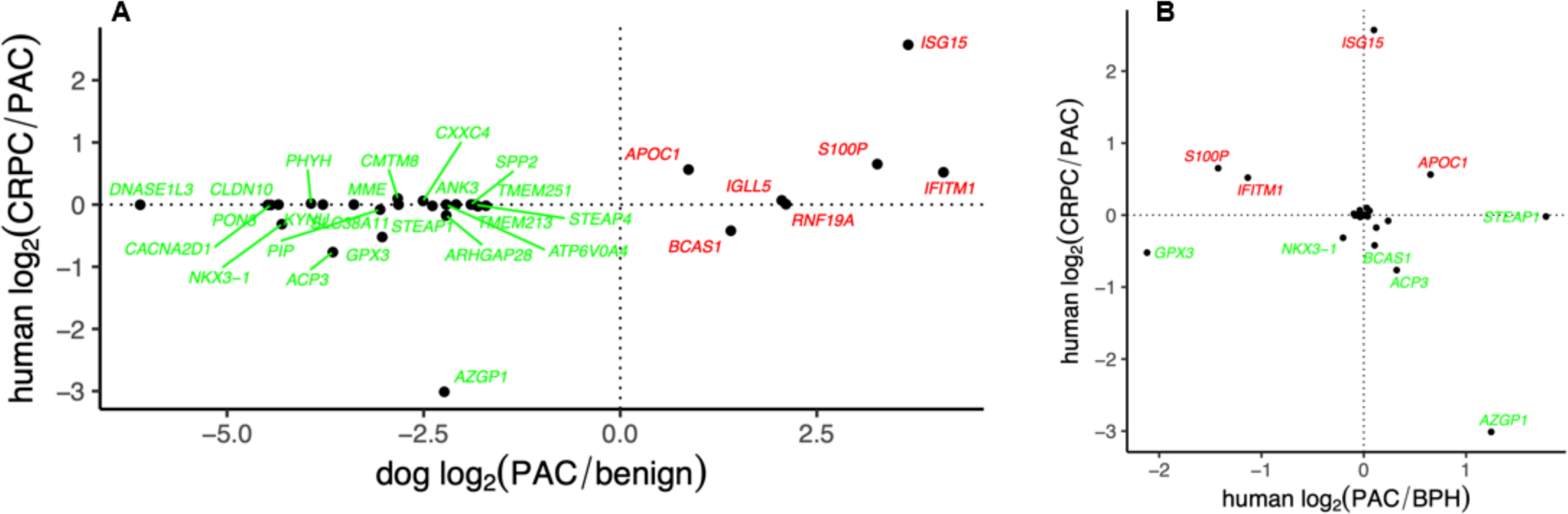
A) Transcript abundance ratios for 33 canine/human ortholog pairs (labeled by official gene symbol) in dogs (prostatic carcinoma vs. benign prostate tissue) and in humans (CRPC vs. localized PC). Two orthologous pairs of genes, *ISG15* and *AZGP1*, showed significant differences in prostate cancer studies in both dogs and humans. **B) Transcript abundance ratios for the genes from the canine prostatic carcinoma analysis, in a human CRPC vs. Localized PC sample group comparison and in a PC vs. BPH sample group comparison.** Among the 33 genes, *ISG15* exhibits greater levels of RNA transcripts in castration-resistant prostate cancer (CRPC) compared to localized prostate cancer. Conversely, *AZGP1* shows lower levels of RNA transcripts in CRPC compared to prostate cancer.

Two orthologous pairs exhibited significant differences in both analyses, consistently exhibiting the same trend. The first pair is ***ISG15*** (ubiquitin-like protein ISG15-like), which is transcriptionally upregulated in prostate cancer in both species. The second pair is ***AZGP1*** (α-2-glycoprotein 1, zinc-binding), which is downregulated in prostate cancer in both species. Other genes that were modestly differentially expressed at the transcript level in human prostate cancer include the upregulated genes *APOC1*, *S100P*, and *IFITM1*, and the downregulated genes *ACP3*, *GPX3*, *NKX3.1*, and *BCAS1*. These results suggested common roles for *ISG15* and *AZGP1* in prostatic carcinoma in dogs and humans, and potential roles for *APOC1*, *S100P*, *IFITM1*, *ACP3*, *GPX3*, *NKX3.1*, and *BCAS1* in both canine and human prostate cancer.

To determine whether the *specificity* of the aforementioned differentially expressed genes’ shift in mRNA abundance in CRPC vs. localized prostate cancer, —i.e., whether the transcript abundance change from localized prostate cancer to CRPC is merely an amplification of an already detectable abundance shift between benign prostate and prostate cancer or (alternately) whether if the shift is unique to castration resistance—we compared transcript abundances of these genes in localized human prostate cancer and in human benign prostate hyperplasia (BPH) using data from the Yun et al.study.^6^ Specifically, we analyzed the 33 genes for their CRPC/localized prostate cancer expression ratios and their localized prostate cancer/BPH expression ratios (Figure 4B). Of the 33 genes, *ISG15* has higher transcript abundance in CRPC than in localized prostate cancer and *AZGP1* has lower transcript abundance in CRPC than in prostate cancer, suggesting that *ISG15* upregulation and *AZGP1* downregulation are specific hallmarks of androgen-indifferent prostate cancer.

Next, we broadened the secondary analysis of published human prostate cancer RNA-seq datasets to obtain transcript abundance ratios for four additional sample group comparisons, for a total of seven sample group comparisons (Table 2; Figure S3).

From a combined view of the five human mRNA abundance ratio types and one canine mRNA abundance ratio type studies, we identified two additional genes that had cross-species mRNA abundance patterns that may suggest a role in CRPC: glutathione peroxidase 3(*GPX3*), which is downregulated in prostatic carcinoma in dogs (vs. benign canine prostate) and downregulated in advanced vs. localized prostate cancer in humans; and S100 calcium binding protein P (*S100P*), which is upregulated in human CRPC vs localized prostate cancer, upregulated in canine prostate cancer vs. benign canine prostate, and downregulated in human prostate cancer vs. human BPH (Figure 5A; Figure S3). Additionally, - interferon-induced transmembrane protein 1-like *IFITM1*; - which had a modestly elevated transcript abundance in human CRPC vs. localized human prostate cancer was markedly elevated in canine prostatic carcinoma vs. canine benign prostate.

**Figure 5.**
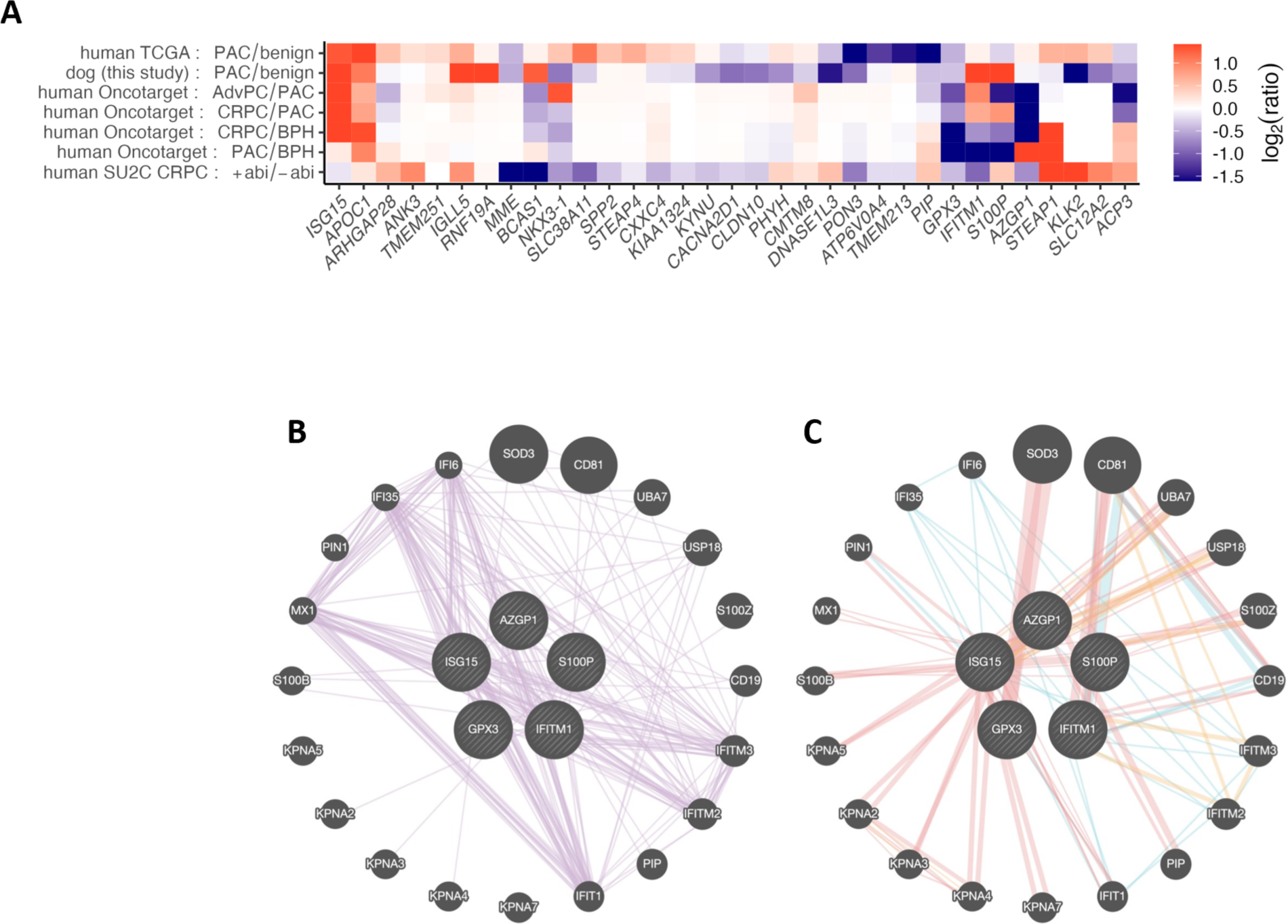
A) mRNA abundance ratios in canine and human prostate cancer studies. Similar patterns in *GPX3* and *S100P*, suggest potential involvement in castration-resistant prostate cancer. *IFITM1* showed slightly higher transcript abundance in human CRPC compared to localized prostate cancer, while its canine counterpart was more abundant in prostatic carcinoma. Rows are organized by similarity of row-wise expression profiles (agglomerative hierarchical linkage). Sample groups labeled as follows: PAC, prostate adenocarcinoma; PAC, prostatic carcinoma; AdvPC, advanced prostate cancer; CRPC, castration-resistant prostate cancer; BPH, benign prostate hyperplasia; +abi, from patient treated with abiraterone; -abi, from patient not previously treated with abiraterone. Transcripts are labeled by official gene symbol. False color indicates log2 relative abundance for the indicated sample group comparison. Columns are ordered to group genes together that have similar column-wise expression patterns. **(B-C). The GeneMania human molecular networks using five seed genes from cross-species prostate cancer transcriptome analysis.** (B) Gene-gene co-expression network. Edges denote significant co-expression at the transcript level across diverse conditions and cell types (based on various transcriptome datasets incorporated in the GeneMania database). (C) Protein-protein interaction network. Edge colors denote interaction type as follows: pink, physical interaction; orange, predicted physical interaction; green, genetic interaction; and cyan, co-participation in a pathway.

In functional genomics, it is well-established that proteins that are co-expressed are more likely to have common functions and participate in common cellular pathways.^35^ Accordingly, we investigated whether the five human genes that were identified in the canine study and in the secondary analysis, *ISG15*, *AZGP1*, *GPX3*, *S100P*, and *IFITM1*, are members of a community of co-expressed genes across various cell types and conditions in the GEO database (Figure 5B).^25^ This analysis yielded a network of 25 genes of which nine genes, *IFIT1*, *IFI6*, *IFI35*, *MX1*, *IFITM2*, *IFITM3*, *USP18*, *ISG15*, and *IFITM,* are type I interferon-induced—this this is an extremely strong enrichment (FDR < 6×10^-12^, odds ratio ∼80) suggesting the involvement of type I interferon signaling in both canine and human prostate cancer. The same community of co-expressed genes has a significant density of interactions or predicted interactions or pathway co-participation relationships among them at the protein-protein level (Figure 5C); out of 300 gene pairs, 29% of gene pairs (i.e., 86 gene pairs) have interactions, 4.8-fold higher than would be expected by chance for random pairs of genes (*p* < 10^-15^, Exact binomial test).

For various types of cancer, gene-gene co-occurrence of somatic mutations is associated with cooperative function or co-participation in specific functional networks.^36,37^ To determine whether this is the case for our gene network, we analyzed the five seed genes for the network analysis (*S100P*, *IFITM1*, *GPX3*, *AZGP1*, and *ISG15*) for co-occurrence of somatic mutations in human prostate cancer. By secondary analysis of human somatic mutation frequency data from the cBioPortal database, we investigated somatic mutation co-occurrence in the five genes.^26^ Importantly, all 5 genes had a highly significant pair-wise co-occurrence of somatic mutations in prostate cancer (Table S2; Figure 6A); in particular, deletions significantly co-occur for *ISG15* and *IFITM1* despite the two genes being on different chromosomes in the human genome, namely, hsa-chr-1 and hsa-chr-11) (Figure 6A). *ISG15* and *IFITM1* are more likely (2.8-fold and 4.7-fold, respectively) to be deleted than amplified in prostate cancer. In contrast, the androgen-responsive gene *AZGP1* is 15-fold more likely to be amplified than deleted (Figure S4). This is consistent with the established androgen sensitivity of *AZGP1*, indicating a possible regulatory function in the androgen signaling cascade.

**Figure 6.**
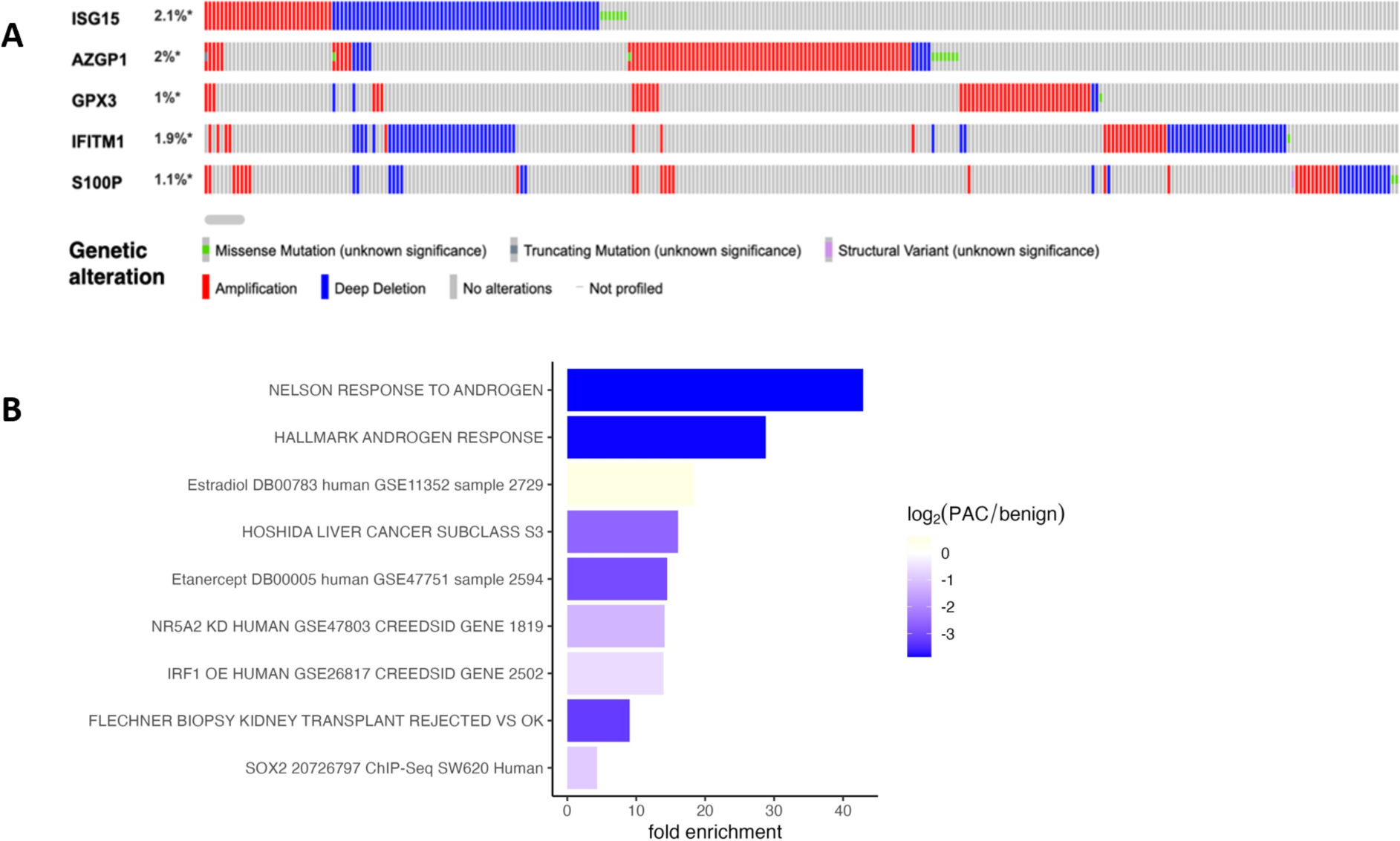
(A) Somatic mutations co-occurrence using cBioPortal database in the five seeded genes for the network analysis. Mutation co-occurrence analysis of *ISG15, AZGP1, GPX3, IFITM1*, and *S100P* across 9,377 prostate tumors. **(B) Enrichment analysis of the 33 differentially expressed genes in canine prostate cancer, for functional gene sets from the Gene Ontology, DrugLib, Enrichr, and MSigDB**. Only gene sets meeting the significance cutoff of FDR < 0.01 are shown. Bar color indicates the average log_2_ expression ratio (PAC/benign) for differentially expressed genes that are in the indicated functional gene set.

Among the 33 genes that are differentially expressed, androgen-responsive genes are enriched 40-fold relative to what would be expected by chance (Figure 6B).

Taken together, these observations suggest that in both spontaneously occurring canine prostatic carcinoma and in human CRPC a group of genes are consistently differentially expressed. Further, multiple lines of evidence, —including co-mutation frequency, co-expression, physical interactions, and functional enrichment —suggest that these genes and their associated network have coordinated function in prostate cancer under low-androgen conditions.

## 4 Discussion

Here, we used mRNA profiles to characterize prostatic carcinoma occurring in castrated dogs, and consequently determine whether these would identify common shared molecular pathways with human castration resistant prostate cancer (CRPC).

Using a microarray approach to analyze RNA from archival FFPE tissues obtained from castrated dogs with prostatic carcinoma, we were able to profile a substantial number of canine genes. Our study focused on 32,391 genes, of which 23,389 were found to be above background in at least one sample, providing a comprehensive view of the transcriptional activity in both prostatic carcinoma and benign canine prostate tissue. By measuring transcript abundances, we identified 33 canine genes with significantly altered transcript abundances in prostatic carcinoma vs. benign prostate (7 upregulated and 26 downregulated).

### 4.1 Androgen-regulated genes and AR signaling

Prostate cancer depends on androgens and AR signaling for growth and progression. In humans, prostate cancer is identified by an increasing level of prostate-specific antigen (PSA), which is AR-regulated. Typically, PSA increases in the setting of human prostate cancer and CRPC.^10^ In contrast, we found that castrated dogs with prostatic carcinoma demonstrated a downregulation of *AR* gene that was validated by the concomitant downregulation of the AR-regulated genes *KLK2*, *NKX31* and *ACP3*, suggesting that prostate cancer occurring in castrated dogs is not driven by *AR*. Other studies have demonstrated similar results in which canine prostatic carcinoma showed progression independently of *AR* and androgen hormone stimulation.^38^ In humans, androgen-indifferent prostate cancer typically develops after treatment with androgen signaling inhibitors in CRPC.^39^ Androgen-indifferent prostate cancer variations include aggressive-variant prostate cancer (AVPC), neuroendocrine prostate cancer (NEPC), and double-negative prostate cancer (DNPC).^40^ Since our findings indicated that *AR* signaling was downregulated in prostate carcinomas arising in castrated dogs, we investigated neuroendocrine differentiation and the genes associated with neuroendocrine prostate cancer (*INSM1, RB1, TP53, CHGB, CHGA*, and *SYPL2*). We did not find any significant changes in the expression of these genes between the canine prostate carcinoma samples and the benign controls (Figure 3).

### 4.2 Comparison of gene expression in human and canine prostatic carcinoma

Our study found two pairs of orthologous genes, *ISG15* and *AZGP1*, that showed persistent and substantial differences in expression levels and directions consistent in both species (Figure 4 A and B). Both humans and dogs with prostate cancer exhibited transcriptional upregulation of *ISG15*. Conversely, prostate cancer in both species resulted in the downregulation of *AZGP1* a zinc-binding alpha-2-glycoprotein.

In human prostate cancer, ***ISG15*** has been previously reported to be transcriptionally over-expressed versus normal prostate tissue and differentially expressed between tumors of patients treated with ADT and those that were not.^41,42^ Further, in the Cancer Genome Atlas (TCGA) prostate adenocarcinoma cohort, high tumoral *ISG15* transcript abundance was found to be associated with reduced survival.^43^

***AZGP1*** has a crucial role in lipid metabolism, glucose utilization, and the control of insulin sensitivity.^44^ Various hormones, most notably androgens, influence the synthesis of *AZGP1.*^44^ Studies have shown that *AZGP1* expression is upregulated during the first phase of prostate cancer progression, but downregulated or entirely missing in advanced prostate cancer.^45^ Additionally multiple studies have identified a correlation between decreased *AZGP1* expression and the occurrence of biochemical failure of patients with prostatic cancer after undergoing radical prostatectomy.^46–48^

In order to determine the specificity of these alterations in gene expression in relation to castration resistance, we compared the levels of transcripts in human CRPC with those in localized human prostate cancer and benign prostatic hyperplasia. Our findings show that increased expression of *ISG15* and decreased expression of *AZGP1* are distinct characteristics of androgen-indifferent prostate cancer, and consequently may be involved in the progression CRPC. The consistency in the direction of expression changes for *ISG15* and *AZGP1* in our study suggests that these genes may play common roles in the development or progression of prostate cancer across species.

Our investigation’s reach was increased by the secondary study of human prostate cancer RNA-Seq datasets, which identified other genes with cross-species mRNA abundance patterns that may be involved in CRPC: ***GPX3*** that plays a crucial function in the process of detoxifying reactive oxygen species (ROS). *GPX3*, which has a tumor suppressor activity and therefore its down-regulation or disruption may contribute to the growth and progression of prostate cancer.^49^ In our study *GPX3* is expressed at lower levels in both canine prostate adenocarcinoma and advanced prostate cancer. ***S100P***, on the other hand, was increased levels in human CRPC and canine prostate cancer. *S100P* is a calcium-binding protein with EF-hand motifs and it is a part of the *S100* family, which acts as regulators of biological processes both within and outside of cells.^49–51^ *S100P* is overexpressed in prostate cancer has been associated with aggressive disease, hormone independent and metastatic phenotype.^52^ ***IFITM1*** (interferon-induced transmembrane protein 1) belongs to a family of cytokines implicated in the innate immune response to viral infections.^53^ *IFITM* proteins have been associated with the majority of cancer characteristics, such as the rapid growth of tumor cells, resistance to treatment, the formation of new blood vessels, the invasion of surrounding tissues, and the spread of cancer to other parts of the body.^54,55^ Correspondingly, in our study there was consistent upregulation of *IFITM1* in both canine PAC and human metastatic CRPC, which implies its role in tumorigenesis and metastases (Figures 4 A and B).

### 4.3 Unveiling co-expression networks and somatic mutation patterns in prostate cancer-associated genes

The integration of functional genomics with the exploration of molecular mechanisms underlying prostate cancer has identified potential roles of co-expressed genes and somatic mutations in disease progression.^36^ Therefore, seeding our analyses with *ISG15*, *AZGP1*, *GPX3*, *S100P*, and *IFITM1* we explored the co-expression networks across different cell kinds by utilizing the GEO database. Consequently, we identified a network of 25 genes connecting these five seed genes. Nine of these genes were identified as type I interferon-induced, with significant functional ∼80-fold enrichment. The network exhibited a striking concentration of connections at the protein-protein level, that the genes within this network are functionally interrelated and may act together in signaling pathways that are disrupted in prostate cancer.

Given the widely accepted idea that the presence of multiple gene mutations within tumors is linked to cooperative function or participation in specific functional networks, we conducted a further analysis of the 5 seed genes to determine if they co-occur with somatic mutations in human prostate cancer.^37^ Consequently, examination of the curated cBioPortal human prostate cancer data revealed highly significant pair-wise co-occurrence of somatic mutations among *S100P*, *IFITM1*, *GPX3*, *AZGP1*, and *ISG15*. Importantly, there was significant co-occurrence of deletions for *ISG15* and *IFITM1*, despite their genomic locations on different chromosomes. These two genes had a notable propensity for deletions in prostate cancer, implying a shared regulatory mechanism in the progression of prostate cancer. The simultaneous presence of somatic mutations and the distinct patterns of deletions and amplifications emphasize the complex genetic makeup of prostate cancer. The observed synergistic functions among these genes, corroborated by both co-expression networks and somatic mutation co-occurrence, underscore the significance of comprehending the wider framework of gene interactions in prostate cancer in the setting of androgen deprivation.

Our cross-species analysis provides a robust framework for identifying genes potentially involved in CRPC. By integrating data from multiple human studies with insights gleaned from canine models, our approach leverages the strengths of both species to uncover genetic pathways that may contribute to CRPC. This dual-species methodology offers a unique perspective, as it allows us to identify common genetic markers and pathways that are conserved across species, thus providing a more comprehensive understanding of the genetic profiling of the pathways involved in castration resistance. Unlike single-species studies, which may be limited by species-specific genetic and environmental factors, our cross-species analysis can highlight evolutionary conserved mechanisms and reveal novel therapeutic targets that might be overlooked when studying only one species. Although it would be beneficial to directly compare castrated and intact dogs with prostatic carcinoma to gain more insights, our current findings serve as a significant initial step in comprehending the genes related with castration-resistant prostate cancer (CRPC) across different species. This integrative approach not only enhances the robustness of our findings but also opens new avenues for translational research, potentially leading to more effective treatments for CRPC.

In summary, our study gives a unique look into the similarities and differences of gene expression between species and identifies a distinct shared molecular profile for castration resistant prostate cancer in both dogs and humans and may inform therapies for androgen indifferent prostate cancer. The identification of consistent patterns across different species provides compelling evidence for generating hypotheses, underscoring the significance of our findings in directing future study.

## Acknowledgements

This project was supported by a Cancer Prevention and Control Team-Building Project grant from Oregon Health & Science University and Oregon State University; the Histopathology Shared Resource for pathology support, the Massively Parallel Sequencing Shared Resource and Integrated Genomics Shared Resource for genomics support (P30 CA069533, and P30 CA069533 13S5 through the OHSU-Knight Cancer Institute); Christina Harrington and Robert Searles for genomics insights and support

