## Supplementary Tables and Figures for "Comparative Transcriptomes of Canine and Human Prostate Cancers Identify Mediators of Castration Resistance"

**Supplementary figures and tables**

Table S1. Summary characteristics of the six dogs whose prostate tumors were profiled in this study. in the study which include breeds, ages, and castration status.

| Breed | Age at death (years) | Castrated |
|---|---|---|
| German Shepard | 9 | Yes |
| Lhasa Apso | 11 | Yes |
| Lhasa Apso | 14 | Yes |
| Border Collie | 11 | Yes |
| Chesapeake Bay Retriever | 13 | Yes |
| Dachshund | 13 | Yes |

**Table S2. Co-occurrence of somatic mutations among the five genes *ISG15*, *AZGP1*, *GPX3*, *IFITM1*, and *S100P*, in human cancers.**

| Gene A | Gene B | Number of cases where neither gene is mutated | Number of cases where gene A is mutated, but not gene B | Number of cases where gene B is mutated, but not gene A | Number of cases where both gene A and B are mutated | Log2 Odds Ratio | p-Value | q-Value |
|---|---|---|---|---|---|---|---|---|
| IFITM1 | ISG15 | 4294 | 53 | 62 | 42 | >3 | <0.001 | <0.001 |
| ISG15 | S100P | 4309 | 89 | 38 | 15 | >3 | <0.001 | <0.001 |
| AZGP1 | GPX3 | 4313 | 88 | 38 | 12 | >3 | <0.001 | <0.001 |
| IFITM1 | S100P | 4315 | 83 | 41 | 12 | >3 | <0.001 | <0.001 |
| AZGP1 | ISG15 | 4262 | 85 | 89 | 15 | >3 | <0.001 | <0.001 |
| GPX3 | S100P | 4355 | 43 | 46 | 7 | >3 | <0.001 | <0.001 |
| AZGP1 | S100P | 4307 | 91 | 44 | 9 | >3 | <0.001 | <0.001 |
| GPX3 | ISG15 | 4305 | 42 | 96 | 8 | >3 | <0.001 | <0.001 |
| AZGP1 | IFITM1 | 4266 | 90 | 85 | 10 | 2.479 | <0.001 | <0.001 |
| GPX3 | IFITM1 | 4312 | 44 | 89 | 6 | 2.724 | <0.001 | 0.001 |

Each row corresponds to a significance analysis for co-occurrence of somatic mutations of the two indicated genes (in columns “Gene A” and “Gene B”), across 4,355 human cancers whose somatic genomes were profiled and the resulting data obtained from the cBioPortal database (see Methods). The four contingency table counts are contained in columns 3–6. The log2 odds ratio is contained in column 7, the *p*-value in column 8, and the FDR-adjusted *p*-value in column 9.

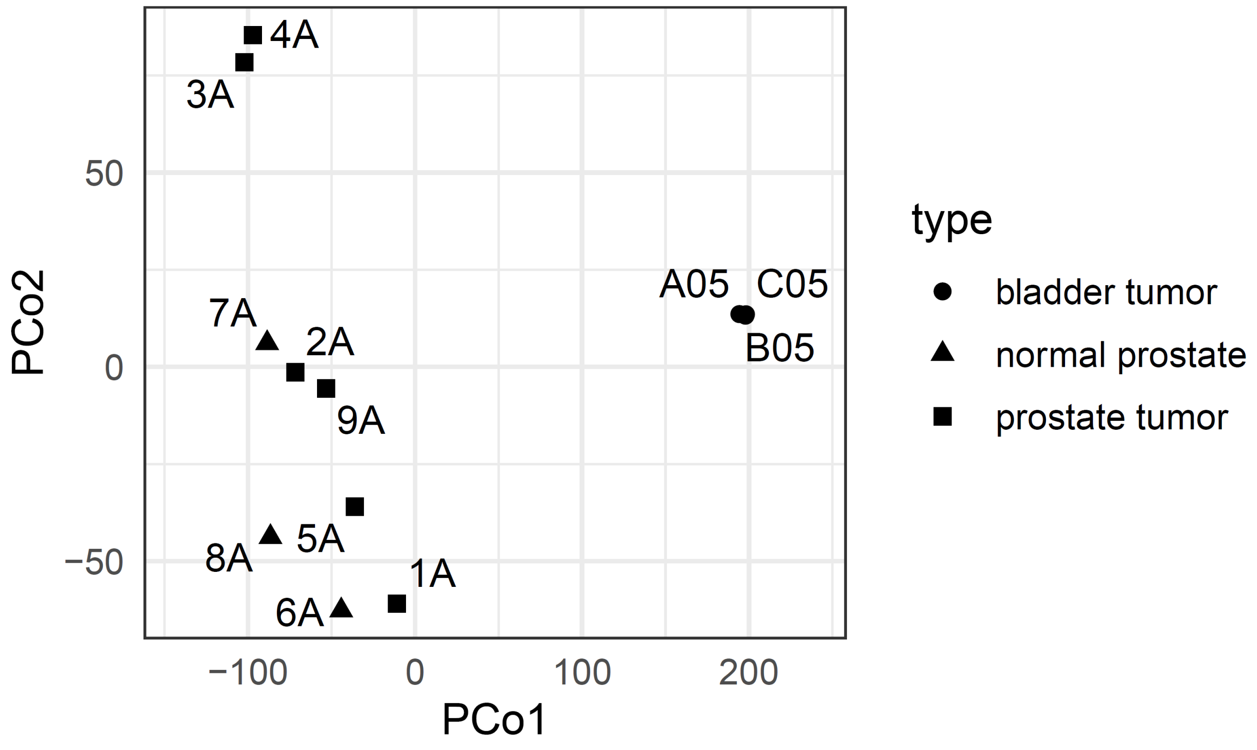


Figure S1: Principal Coordinate Analysis (PCoA) of Bladder and Prostate Samples

PCoA plot was generated using normalized data (see Methods) and demonstrates clear separation between bladder and prostate tumor samples along PCo1, with further distinction between normal and tumor prostate samples primarily along PCo2. This visualization suggests significant molecular differences between tissue types, and between normal and tumor states in prostate tissue.

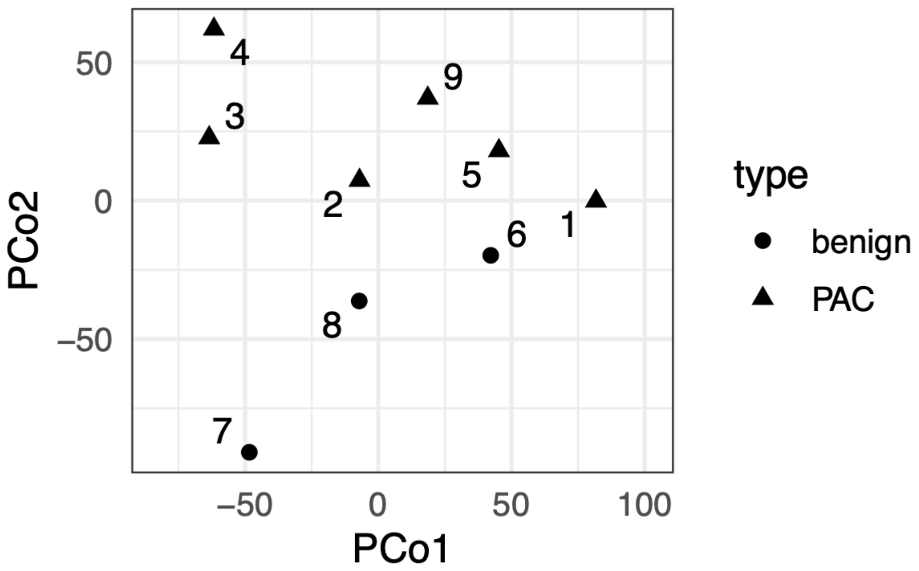


Figure S2. Principal Coordinate 2 separates canine normal prostate samples from prostate adenocarcinoma samples. Each mark represents a tissue sample (sample group indicated by shape). Marks are positioned on the chart using the Principal Coordinates Analysis (PCoA) algorithm (see Methods), in which positions are assigned so that for each pair of samples, the distance between them on the chart most closely correlates with the difference between the samples' transcriptomes.

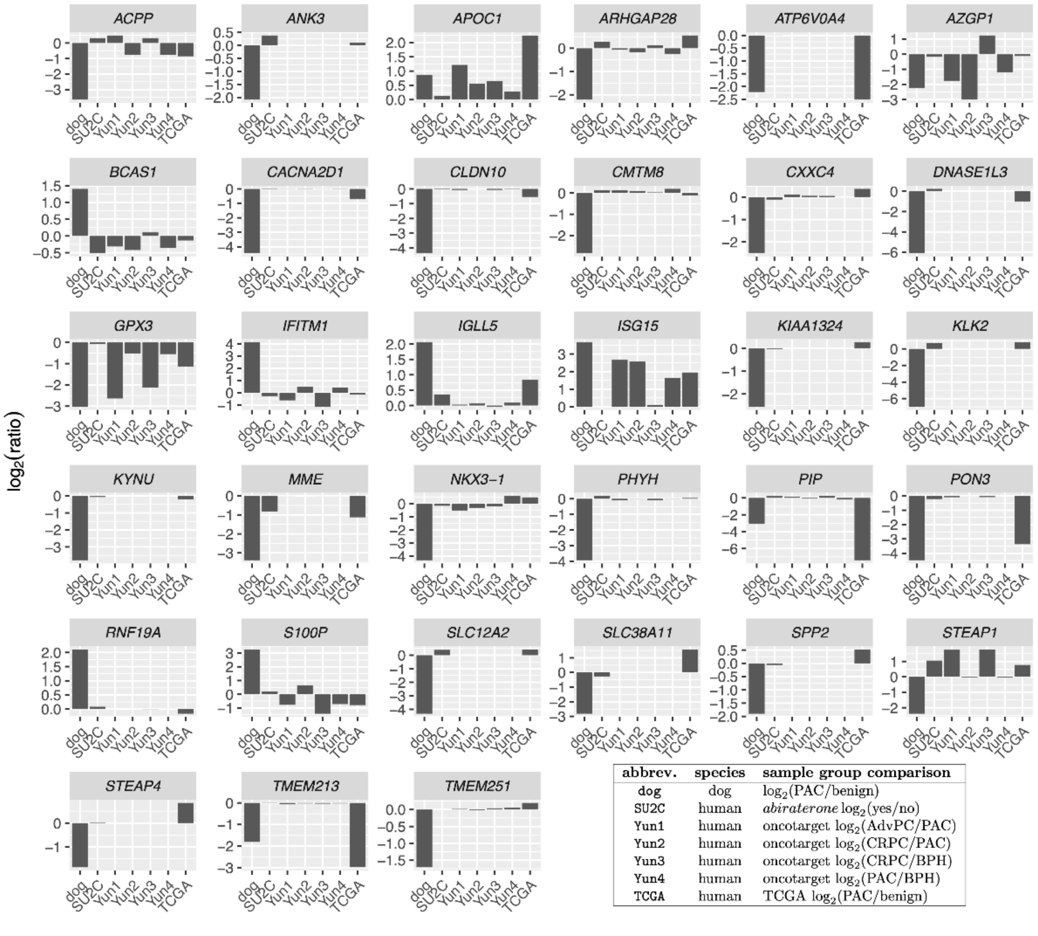


Figure S3. Faceted plot of 33 gene-level transcript abundance $\log_2$ ratios in seven different sample group comparisons (see legend). Each bar chart corresponds to a specific gene (see chart titles) for which orthologs could be mapped between canine and human and for which there was differential expression between benign and prostate adenocarcinoma in dog. Each of the seven types of $\log_2$ ratios is for a specific sample group comparison in our dog study or in a human prostate cancer study (see Methods).

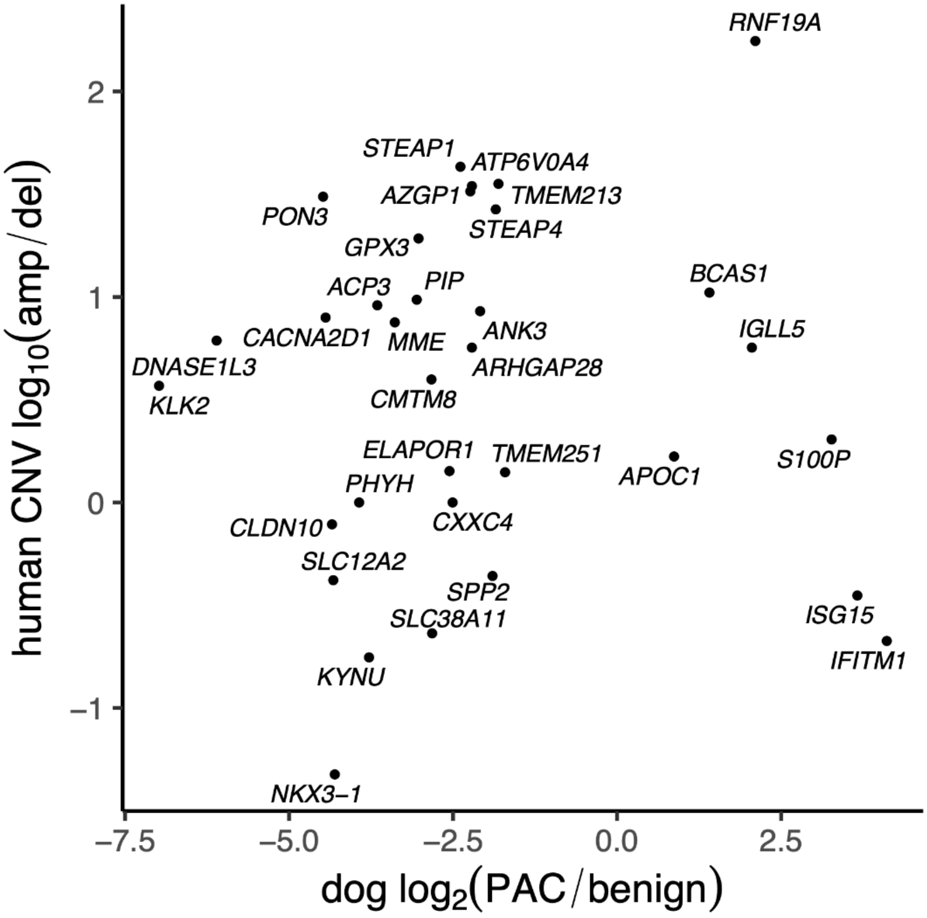


Figure S4. Comparison of human somatic copy number variation and canine transcript abundance ratios, for 33 genes. Each mark corresponds to a specific gene (see mark labels) for which orthologs could be mapped between canine and human and for which there was differential expression between benign and prostate adenocarcinoma in dog. The vertical axis measures the $\log_{10}$ of the ratio (in human prostate cancer) of the frequencies of somatic copy number amplifications to deletions, for the indicated gene (see Methods).
